# Elevated LRRK2 and α-synuclein levels in CSF of infectious meningitis patients

**DOI:** 10.1101/599381

**Authors:** Susanne Herbst, Suzaan Marais, Maximiliano G. Gutierrez, Simon J Waddell, Robert J. Wilkinson, Rachel PJ Lai

## Abstract

Neurodegenerative diseases such as Parkinson’s (PD) have a complex aetiology consisting of an interplay of genetic and environmental factors. Inflammation and infection are proposed external factors that trigger disease progression. Tuberculous and cryptococcal meningitis frequently lead to long-term neurological sequelae but their association with the development of PD are unexplored. In this study, we protein profiled the CSF from 76 patients with or without infectious meningitis and found that proteins commonly associated with PD (LRRK2, tau and alpha-synuclein) were significantly elevated, establishing a link between neuroinflammation and infection. Importantly, these findings suggest that LRRK2, tau and alpha-synuclein could represent biomarkers of neuroinflammation.

## Introduction

An increasing body of evidence implicates neuroinflammation in the aetiology of neurodegenerative diseases such as Parkinson’s Disease (PD). It has been speculated that infection-induced inflammation can lead to damage in neuronal viability and functionality, thus contributing to neurodegenerative disease progression^1^. Elevated concentrations of pro-inflammatory cytokines have been detected in both cerebrospinal fluid (CSF) and post-mortem brain tissue samples of PD patients^2^. The precise aetiology of PD is largely unknown and likely consists of a complex interplay between genetic and environmental factors. The most common underlying genetic cause for inherited forms of PD are mutations in Leucine-rich repeat kinase 2 (LRRK2) and it has been hypothesised that infectious diseases constitute an environmental trigger for PD development^3^. Additionally, infections in the periphery are known to worsen motor function in PD patients, indicating that inflammatory mediators directly impact disease progression^4^.

Infectious meningitis caused by either *M. tuberculosis* or *C. neoformans* is characterised by the increase of inflammatory and neural injury makers in the meninges^5-8^. In both tuberculous and cryptococcal meningitis (TBM and CM), up to half of survivors experience neurological sequelae with deficits that may be similar to those in neurodegenerative diseases, such as impaired cognition and movement disorder.

Here we report the results of a large-scale proteomic analysis comparing protein signatures in the CSF from patients with or without infectious meningitis (TBM, CM, or viral meningoencephalitis, VM). We identified a cluster of proteins that is functionally associated with neurodegenerative diseases, including LRRK2, α-synuclein and tau. CSF abundance of these proteins was elevated in patients with TBM and CM and positively correlated with inflammatory cytokines. Together, the data suggest that neurodegeneration-associated proteins can be considered inflammatory markers themselves that respond to infectious disease triggers.

## Methods

Adults (age ≥18) with suspected meningitis who underwent lumbar puncture as part of their diagnostic workup were recruited into a diagnostic study^9^ at Mitchell’s Plain Hospital and Khyelitsha Hospital, Cape Town, South Africa. Patients were excluded if bacterial meningitis other than TB was suspected (cloudy or pus-like CSF). The study was approved by the University of Cape Town Human Research Ethics Committee (HREC REF: 730/2014). Informed consent was obtained from all fully conscious patients. Patients with impaired consciousness were enrolled and patient consent was sought when capacity was regained. If death occurred before capacity was regained data was included following ethical approval.

TBM was diagnosed using the consensus case definition^10^ where (i) definite cases had at least one of the following CSF findings: acid-fast bacilli seen, *Mtb* cultured or GeneXpert MTB/RIF positive and (ii) probable cases had a total diagnostic score of ≥12 (if cerebral imaging was performed) or ≥10 (if no cerebral imaging was performed). Possible TBM cases and possible pyogenic meningitis cases were excluded from this study. CM was diagnosed by positive CSF cryptococcal latex antigen test or culture. Patients were classified as VM if they presented with symptoms and signs of meningitis, had raised CSF lymphocytes (with or without raised protein and decreased glucose) and recovered without treatment directed at any specific organism. Herpes simplex virus (HSV) meningitis was diagnosed by positive CSF HSV PCR or by a good clinical response to acyclovir. Patients without CNS infection (other than HIV-1) who presented with chronic headaches, psychosis, or HIV-associated neurocognitive disorder were included as non-meningitis controls. HIV status was known for all participants.

SOMAscan, an aptamer-based multiplexed proteomics assay, was used to measure the abundance of protein analytes in CSF samples (SomaLogic, Inc.; Boulder CO, USA). Briefly, SOMAmer reagents bind with high affinity and specificity to their cognate protein target in the CSF, which are then released and hybridised to a DNA array, resulting in relative luminescence units (RLU) as a readout that is directly proportional to the concentration of the corresponding protein in each CSF sample. SOMAscan data of all samples were hybridisation-normalised and adjusted for plate scaling factors calculated from signals from the control probes. Statistical differences between patient groups of each protein of interest were calculated using two-tailed Mann Whitney *U* test and *p*<0.05 was considered significant. Correlation analysis was performed using the ggpubr package on R using Spearman’s Rank and *p*<0.05 was considered significant.

For Western Blotting, CSF proteins were precipitated with methanol (1:3 v/v). Proteins were detected with the following antibodies: LRRK2 (clone N241A/34, NeuroMab; Davis CA, USA), α-synuclein (clone MJFR1, Abcam, Cambridge UK), Tau (clone D1M9X) and β-actin-HRP (both from Cell Signaling; Hitchin UK). Protein loading was assessed by Ponceau S staining.

## Results

A total of 76 patients were enrolled into this study, including 20 with TBM (13 definite and 7 probable; HIV-infected=17); 24 with CM (HIV-infected=24); seven with VM (HIV-infected=3) of whom two had HSV meningitis; and 25 controls without meningitis (HIV-infected=6).

As previously reported, the proteomic analysis of CSF identified increased levels of inflammatory cytokines, such as TNF-α, IL-1β and IFN-β in all meningitis samples (**Fig 1A**). This was paralleled by an increase in cerebral injury markers such as matrix metallopeptidase 9 (MMP-9), glial fibrillary acidic protein (GFAP) and ubiquitin C-terminal hydrolase L1 (UCH-L1) (**Fig 1B**). TBM induced the highest increase in both inflammatory cytokines and injury markers, and VM consistently showed only a mild increase of these proteins. In addition to the inflammatory protein pattern, we observed an unexpected increase in a group of proteins typically associated with neurodegenerative diseases, such as LRRK2, tau and α-synuclein in patients with CM and TBM. Specifically, TBM showed 2-fold higher median levels in tau and α-synuclein and a striking 10-fold elevation in LRRK2 levels, when compared to non-meningitis controls (**Fig 2A**). We validated the proteomics findings by Western Blotting and confirmed that LRRK2, α-synuclein and tau were significantly elevated in TBM, and to a lesser extent in CM, when compared to CSF from control individuals (**Fig 2B**).

**Figure 1:**
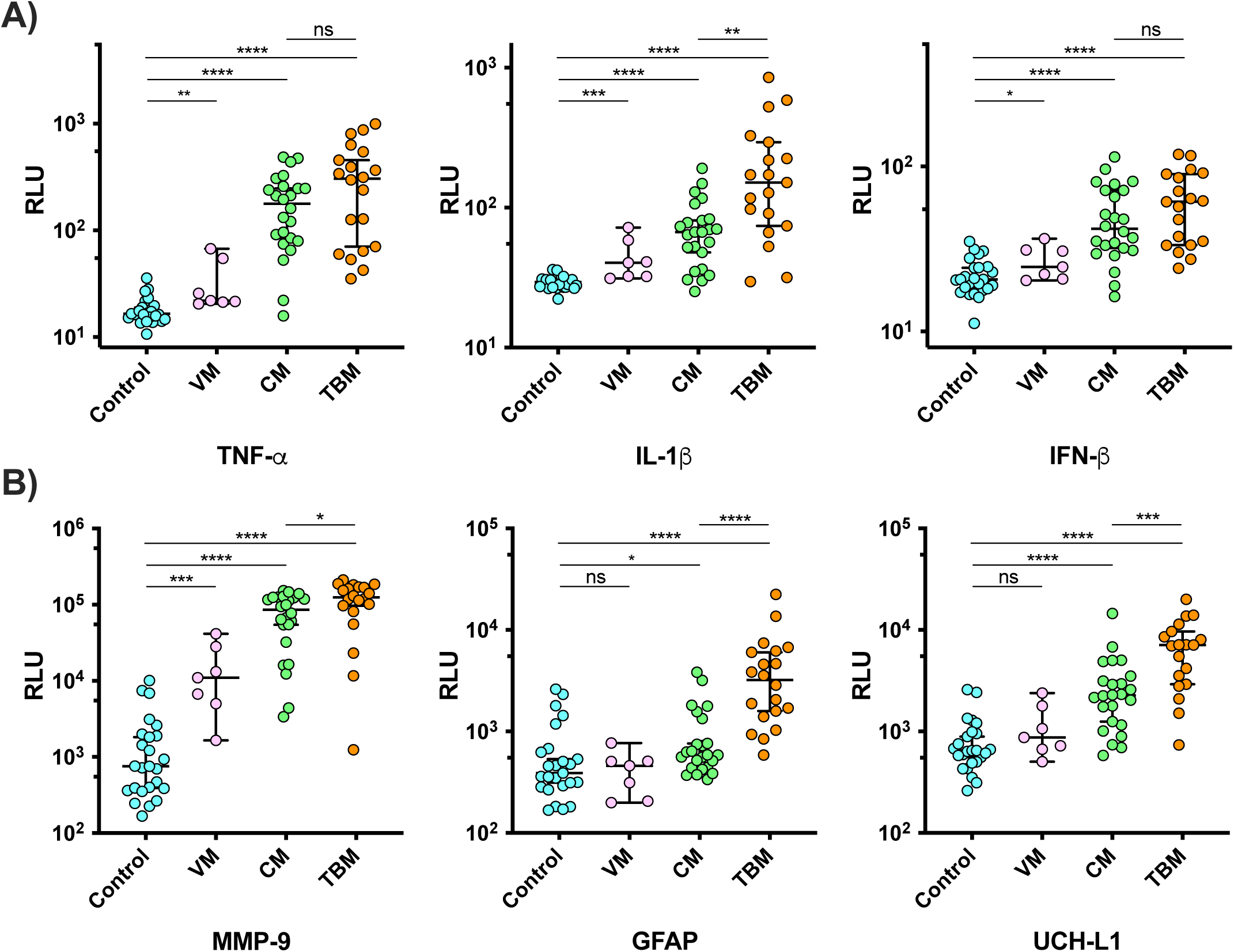
Cerebrospinal fluid abundance of (**A**) inflammatory markers and (**B**) brain injury markers were measured in control patients without meningitis and in those suffering from viral (VM), cryptococcal (CM) and tuberculous (TBM) meningitis.

**Figure 2:**
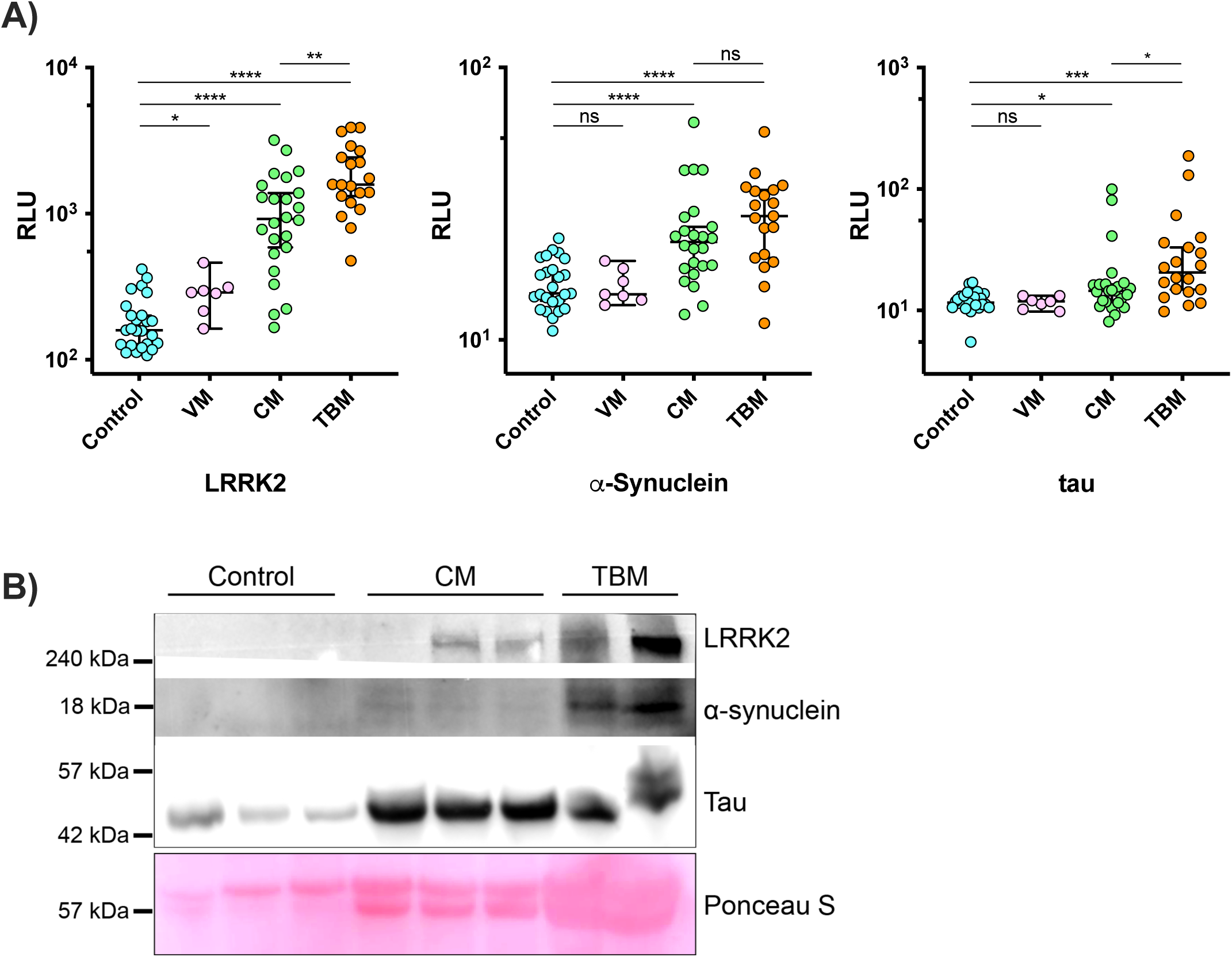
**(A)** Cerebrospinal fluid abundance of neurodegeneration-associated proteins were measured in control patients without meningitis and in those with viral (VM), cryptococcal (CM) and tuberculous (TBM) meningitis. **(B)** Western Blotting of CSF proteins for LRRK2, Tau and α-synuclein. Ponceau S stain is used as an indicator for total protein amounts.

Sub-analyses based on outcome were performed on CM and TBM where a subset of patients died, thus were considered to have more severe infection at the time of sampling. While disease severity impacted the CSF abundance of inflammatory cytokines (**Figure 3A**), there were no difference in the levels of these neurodegenerative disease-associated proteins (**Figure 3B**), suggesting that the aetiology may be a more important determinant. Sub-analysis on the impact of HIV-1 coinfection was not performed on the three meningitis groups as the number of non-HIV-1 infected patients was too small for statistical analysis.

**Figure 3:**
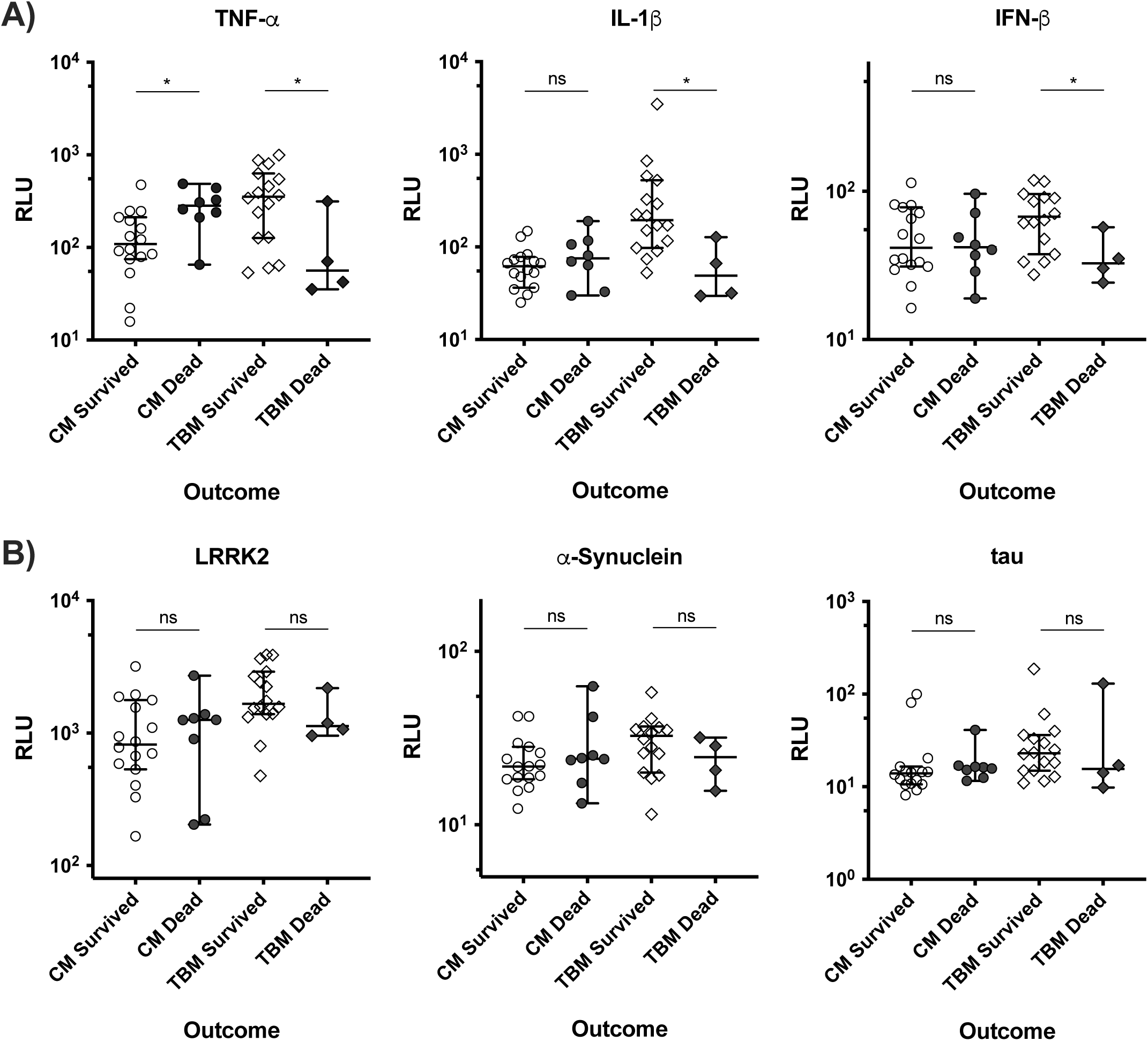
Sub-analyses were performed to decipher the impact of disease severity on cerebrospinal fluid abundance of (**A**) inflammatory cytokines and (**B**) neurodegeneration-associated proteins in patients with cryptococcal (CM) and tuberculous (TBM) meningitis. There was no mortality in control non-meningitis patients or those with viral meningitis.

Studies in PD showed that the CSF levels of neurodegeneration-related proteins significantly correlate with inflammatory cytokines including TNF-α and IL-6^11,12^. Similarly, we found a significant correlation between LRRK2 and the inflammatory cytokines TNF-α and IL-1β and the neural injury marker UCH-L1 in TBM, CM and VM (**Fig 4**) suggesting that LRRK2 expression is associated with inflammatory responses.

**Figure 4:**
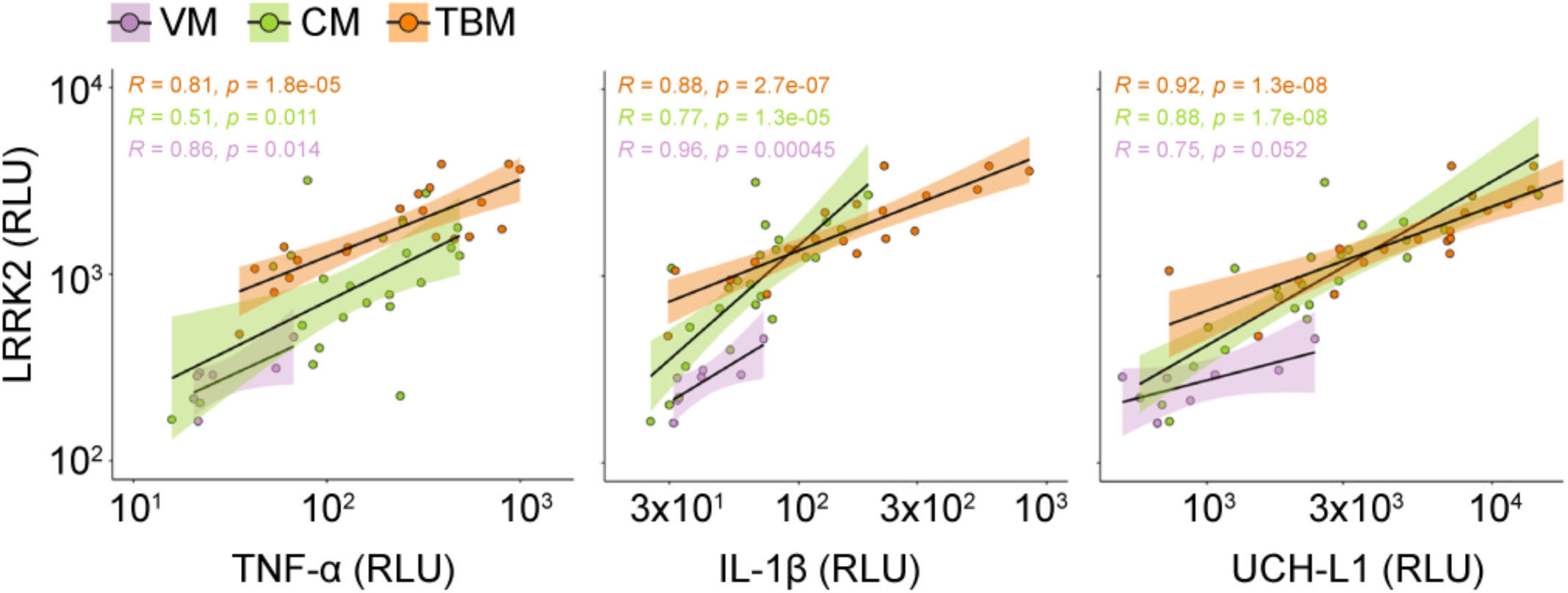
Correlation analysis between LRRK2 and the inflammatory mediators (**A**) TNF-α and (**B**) IL-1β and (**C**) the brain injury marker UCH-L1 in CSF samples of patients suffering from viral meningitis (VM), cryptococcal meningitis (CM) and tuberculous meningitis (TBM).

## Discussion

In this study, we show a significant and selective upregulation of proteins associated with neurodegenerative diseases in the CSF of patients suffering from tuberculous and cryptococcal meningitis. Our findings implicate these proteins as potential markers of neuroinflammation and/or brain damage that could functionally contribute to meningitis pathology. The up-regulation of LRRK2 in the CSF additionally raises the question if infections can affect long-term brain function by resulting in the infiltration of LRRK2 and α-synuclein expressing monocytes/neutrophils from the periphery, thereby contributing to PD development in genetically susceptible individuals. This is of special interest as the penetrance of LRRK2 mutations is incomplete, ranging from 40-75% at the age of 80^13,14^. As such, our findings indicate infections might constitute an environmental factor that contributes to disease development.

Studies examining the history of CNS infections in a general population of PD patients have found only a weak correlation between PD disease development and CNS infections^15^. Importantly, longitudinal studies that specifically examine the relationship between infectious disease history and PD development in LRRK2 mutation carriers are lacking. It has been shown that α-synuclein is also up-regulated in the enteric nervous system during inflammation or after viral infection^16^ and our results provide additional evidence supporting the idea that α-synuclein is implicated in inflammatory processes. Several studies explored the monitoring of LRRK2 expression levels in peripheral blood or CSF as a potential biomarker for PD^17,18^. Here, we found that in CSF of patients with infectious meningitis, LRRK2 protein levels are significantly increased and robustly correlated with the protein levels of the inflammatory cytokines TNF-α or IL-1β. The findings reported here provide further evidence for an association between LRRK2 with inflammation, immunity and infection. We only observed a significant increase of LRRK2, α-synuclein and tau in patients suffering from tuberculous and cryptococcal meningitis but not viral meningitis. We cannot definitively conclude that specific infectious agents cause an upregulation of this protein cluster in the CSF, as the VM patients only showed a mild increase in inflammatory and injury markers when compared to CM and TBM patients indicating that disease severity might be a determining factor. Nevertheless, in CM and TBM disease severity, as defined by outcome, is not associated with the elevation of these neurodegeneration-related proteins.

Overall, this proteomic analysis suggest that certain proteins associated with neurodegeneration, and PD in particular, can be considered as markers of inflammation, and that inflammatory triggers such as infectious diseases can result in the shuttling of these neurotoxic proteins to the CNS.

## Acknowledgements

The authors would like to acknowledge the contributions of the laboratory staff at the Center for Infectious Diseases Research in Africa (CIDRI-Africa) and the doctors and nurses who cared for the study patients and the patients who participated. We would also like to thank Muki Shey for assisting with sample preparation. SH, MG and RJW were funded by the Francis Crick Institute which receives core funding from Cancer Research UK, the UK Medical Research Council and the Wellcome Trust (FC001092 to MGG and FC00110218 to RJW). RJW and SM are supported by the Wellcome Trust (104803 and 203135 for RJW and 097254 for SM). RPJL is supported by the UK Medical Research Council (MR/R008922/1).

## Author Contributions

SM, RJW and RPJL conceived and designed the study. SM recruited, sampled and collected data from patients. SH, SM and RPJL performed experiments and analysed the data. SW provided data acquisition tool. SH, SM, MGG, RJW and RPJL wrote the manuscript with inputs from SW.

## References

1. Perry, V.H., Cunningham, C. & Holmes, C. Systemic infections and inflammation affect chronic neurodegeneration. Nat Rev Immunol 7, 161–167 (2007).

2. Lindqvist, D., et al. Cerebrospinal fluid inflammatory markers in Parkinson’s disease-associations with depression, fatigue, and cognitive impairment. Brain Behav Immun 33, 183– 189 (2013).

3. Johnson, M.E., Stecher, B., Labrie, V., Brundin, L. & Brundin, P. Triggers, Facilitators, and Aggravators: Redefining Parkinson’s Disease Pathogenesis. Trends Neurosci 42, 4–13 (2019).

4. Brugger, F., et al. Why is there motor deterioration in Parkinson’s disease during systemic infections-a hypothetical view. NPJ Parkinsons Dis 1, 15014 (2015).

5. Marais, S., et al. Neutrophil-associated central nervous system inflammation in tuberculous meningitis immune reconstitution inflammatory syndrome. Clin Infect Dis 59, 1638–1647 (2014).

6. Rohlwink, U.K., et al. Biomarkers of Cerebral Injury and Inflammation in Pediatric Tuberculous Meningitis. Clin Infect Dis 65, 1298–1307 (2017).

7. Lan, S.H., Chang, W.N., Lu, C.H., Lui, C.C. & Chang, H.W. Cerebral infarction in chronic meningitis: a comparison of tuberculous meningitis and cryptococcal meningitis. QJM 94, 247–253 (2001).

8. Tugume, L., et al. HIV-Associated Cryptococcal Meningitis Occurring at Relatively Higher CD4 Counts. J Infect Dis 219, 877–883 (2019).

9. Heemskerk, A.D., et al. Improving the microbiological diagnosis of tuberculous meningitis: A prospective, international, multicentre comparison of conventional and modified Ziehl-Neelsen stain, GeneXpert, and culture of cerebrospinal fluid. J Infect (2018).

10. Marais, S., et al. Tuberculous meningitis: a uniform case definition for use in clinical research. Lancet Infect Dis 10, 803–812 (2010).

11. Delgado-Alvarado, M., et al. Tau/alpha-synuclein ratio and inflammatory proteins in Parkinson’s disease: An exploratory study. Mov Disord 32, 1066–1073 (2017).

12. Hall, S., et al. Cerebrospinal fluid concentrations of inflammatory markers in Parkinson’s disease and atypical parkinsonian disorders. Sci Rep 8, 13276 (2018).

13. Ferreira, M. & Massano, J. An updated review of Parkinson’s disease genetics and clinicopathological correlations. Acta Neurol Scand 135, 273–284 (2017).

14. Lee, A.J., et al. Penetrance estimate of LRRK2 p.G2019S mutation in individuals of non-Ashkenazi Jewish ancestry. Mov Disord 32, 1432–1438 (2017).

15. Fang, F., et al. CNS infections, sepsis and risk of Parkinson’s disease. Int J Epidemiol 41, 1042–1049 (2012).

16. Stolzenberg, E., et al. A Role for Neuronal Alpha-Synuclein in Gastrointestinal Immunity. J Innate Immun 9, 456–463 (2017).

17. Cook, D.A., et al. LRRK2 levels in immune cells are increased in Parkinson’s disease. NPJ Parkinsons Dis 3, 11 (2017).

18. Wang, S., et al. Elevated LRRK2 autophosphorylation in brain-derived and peripheral exosomes in LRRK2 mutation carriers. Acta Neuropathol Commun 5, 86 (2017).

